# A novel pH-sensitive reporter reveals the cocaine-regulated trafficking of dopamine transporters in neuronal processes

**DOI:** 10.1101/2022.10.12.511970

**Authors:** Jacqueline Saenz, Oscar Yao, Meha Aggarwal, Xiaofeng Zhou, David J. Barker, Emanuel DiCicco-Bloom, Ping-Yue Pan

## Abstract

The dopamine transporter (DAT) mediated DA reuptake is a major molecular mechanism for termination of dopaminergic signaling in the brain. Psychoactive substances such as cocaine act by inhibition of plasma membrane DAT function as well as by altering its expression. The precise manner and mechanism by which cocaine regulates DAT trafficking, especially at neuronal processes, are poorly understood. We have now engineered a novel pH-sensitive reporter for DAT by conjugating pHluorin to the second exofacial loop of human DAT. We show that DAT-pHluorin can be used to study DAT localization and its dynamic trafficking at neuronal processes. Using DAT-pHluorin we show that unlike neuronal soma and dendrites, which contain majority of the DATs in weakly acidic intracellular compartments, axonal DATs at both shafts and boutons are primarily (75%) localized to the plasma membrane, while varicosities contain abundant intracellular DAT within acidic intracellular structures. Using this novel reporter, we show, for the first time, that cocaine exposure leads to a brief DAT internalization followed by membrane reinsertion that lasts for days. We further show that the cocaine-induced DAT trafficking is sensitive to the activities of Synaptojanin1 phosphatase. Thus, our study using the newly engineered DAT optical reporter reveals the previously unknown dynamics and molecular regulation for cocaine-regulated DAT trafficking in neuronal processes.

## Introduction

Dopamine (DA) signaling regulates a multitude of brain functions including motor coordination and reward processing. DA reuptake by the plasma membrane dopamine transporter (DAT) from the extracellular space into the cytosol is the primary mechanism for termination of brain DA signaling. Lack of DAT expression results in increased extracellular DA and hyperactive locomotion in the knockout mice (Jones et al. 1998). Importantly, many psychoactive substances, such as cocaine and amphetamine (AMPH), directly or indirectly target plasma membrane DAT. In addition to acting to increase extracellular levels of DA, these substances are also known to impact DAT trafficking and plasma membrane expression (Kahlig and Galli 2003; Zahniser and Doolen 2001). The potency of these psychoactive substances in altering DA signaling is highly dependent on the amount of surface DAT expression, however, our understanding on the dynamic trafficking and molecular regulation of DAT, especially in neuronal processes, is poorly understood.

The lack of understanding largely reflects a lack of molecular tools to examine the dynamic trafficking of DAT in small neuronal structures like the axons and presynaptic terminals. Current understanding of DAT trafficking is largely based on studies of heterologous cells and isolated synaptosomes. The most commonly used research strategies to examine the surface DAT include 1) biotinylation assays, which has poor temporal resolution; 2) radioactive binding/uptake assays, which entangles DAT localization with function; and 3) imaging studies using fluorescent proteins-tagged DATs or fluorescent DAT ligands (Eriksen et al. 2009; Guthrie et al. 2020), which exhibit limited spatial resolution for neuronal processes. Complex regulatory mechanisms have been proposed for DAT in different cell systems using different methods. For example, the constitutive DAT recycling can be regulated through a protein kinase C (PKC)-dependent (Loder and Melikian 2003) or independent manner (Sorkina et al. 2005). AMPH was shown in most studies to induce DAT internalization via RhoA and TAAR1 signaling (Wheeler et al. 2015; Boudanova, Navaroli, and Melikian 2008; Underhill et al. 2021), however, AMPH-induced delivery of DAT to the plasma membrane has also been reported (Johnson et al. 2005). Cocaine was found in various studies to block DAT internalization and to increase its surface expression (Saunders et al. 2000; Little et al. 2002; Daws et al. 2002; Underhill et al. 2014; Siciliano et al. 2018; Sorkina et al. 2021). Interestingly, however, the mechanisms underlying cocaine-regulated DAT trafficking remains elusive, likely due to its inhibitory role on DAT function that hinders the use of many conventional methods. An earlier study by Sorkin and colleagues further suggested that neuronal DAT trafficking might be regulated by different mechanisms than what we learned from heterologous cells (Block et al. 2015).

To better reveal and understand DAT trafficking in neurons, we sought to develop a genetically-encoded optical reporter. The pH-sensitive “pHluorin” is a variant of green fluorescent proteins (GFP) that fluoresces upon deprotonation (Miesenbock, De Angelis, and Rothman 1998). PHluorin-based assays are best known for analyzing synaptic vesicle recycling when pHluorin is tagged to synaptic vesicle proteins, such as vesicle associated membrane protein 2/VAMP2 (synaptopHluorin) (Miesenbock, De Angelis, and Rothman 1998; Sankaranarayanan et al. 2000), vesicular glutamate transporter 1/vGLUT1 (Pan, Marrs, and Ryan 2015; Voglmaier et al. 2006), and vesicular monoamine transporter 2/vMAT2 (Pan et al. 2017; Pan and Ryan 2012). Additionally, pHluorin has also been successfully engineered into plasma membrane cargo proteins, such as AMPA receptors (Araki, Lin, and Huganir 2010; Graves et al. 2021), transferrin receptors (Merrifield, Perrais, and Zenisek 2005), mu-opioid receptors (Jullie et al. 2020; Yu et al. 2010) and glucose transporter 4 (GluT4) (Ashrafi et al.

2017; Burchfield et al. 2013) for analyzing their endosomal trafficking. We have now engineered pHluorin into the second exofacial loop of human DAT. By expressing DAT-pHluorin in cultured midbrain (MB) neurons, we turn DAT trafficking events at nano/micrometer scales to measurable changes in fluorescence using a conventional wide-field microscope. We demonstrate that 75% of axonal DAT is on the plasma membrane, with most intracellular DAT found in the varicosities. Moreover, we reveal the dynamic trafficking of DAT to the axonal surface in response to cocaine, which lasts for up to 4 days. This process is regulated by synaptojanin1, a phosphoinositol phosphatase that was previously known to regulate synaptic vesicle recycling (McPherson et al. 1996; Cremona et al. 1999; Verstreken et al. 2003; Mani et al. 2007; Dong et al. 2015) and autophagy (Vanhauwaert et al. 2017; Yang et al. 2022). Thus, our study using the newly engineered DAT optical reporter reveals, for the first time, cocaine-regulated DAT trafficking in neuronal processes.

## Results

### Engineering and characterization of a pH-sensitive DAT reporter

DAT belongs to the family of Na^+^/Cl^-^ dependent neurotransmitter transporters (Amara and Kuhar 1993) and contains 12 transmembrane domains. An extensive body of work has shown that DAT substrates, such as DA, AMPH or cocaine, can induces DAT trafficking (Kahlig and Galli 2003; Zahniser and Doolen 2001). To engineer a DAT reporter suited for reporting physiologically relevant DAT endosomal trafficking, our strategy is to insert pHluorin while not, or minimally, disrupting its endogenous functions in substrate binding. Among the six extracellular loops of human DAT cDNA, the second loop was selected due to the optimal length of the loop that provides flexibility for accommodating a tag protein (**Fig. 1A**). Flexible linkers were inserted on both ends of pHluorin to introduce additional flexibility for the proper folding of the reporter. PHluorin was inserted at multiple sites, T173/E174, P194/G195, G199/S201, G209/T210, R219/G220 outside the known functional sites of DAT (Beuming et al. 2008; Li et al. 2004; Nielsen et al. 2019; Sorkina et al. 2009). While some insertion sites (T173/E174, G209/T210) resulted in poor expression of the transgene in the initial screen, little difference was observed for the remaining versions of DAT-pHluorin (DAT-pH) in their overall expression patterns (data not shown). We thus proceeded to further validate the DAT-pH at the P194/G195 insertion site, where an HA epitope was inserted in earlier studies (Rao et al. 2012; Sorkina et al. 2006).

**Figure 1.**
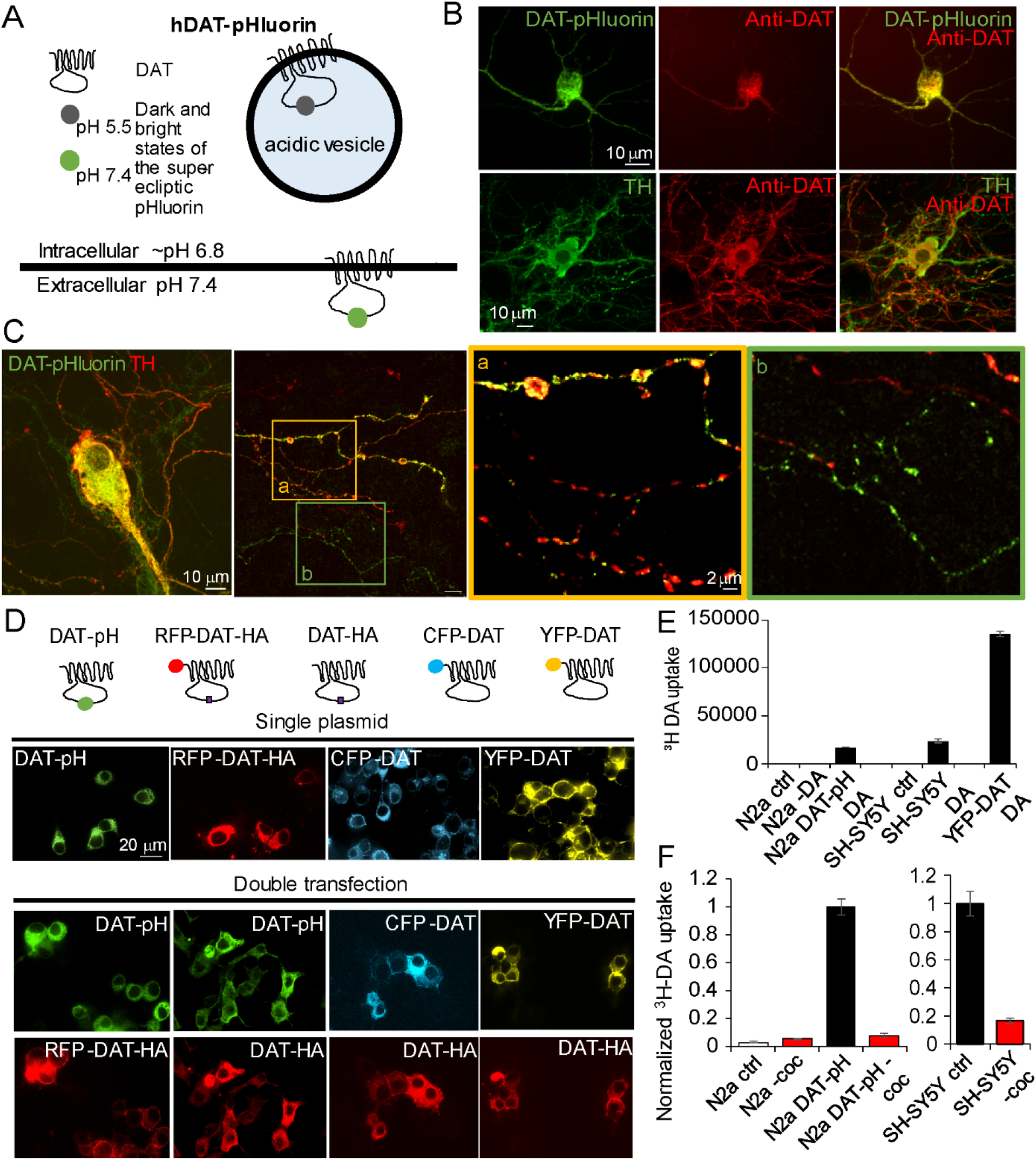
Engineering of a pH-sensitive DAT reporter. (A) Illustration for the engineering of the pH-sensitive DAT reporter. The pH-sensitive GFP variant, pHluorin, was inserted between the 194 Proline and 195 Glycine of the second exofacial loop of human DAT. This design allows DAT-pHluorin to fluoresce at its brightest when inserted onto the plasma membrane to face a neutral (pH 7.4) extracellular medium; and quenches when endocytosed to face an acidic endocytic compartment. (B) Top, cultured MB neuron expressing DAT-pH and immunolabeled with anti-GFP (green) and anti-DAT antibodies (Millipore MAB369) (red). Bottom, cultured MB neuron immunolabeled with anti-TH (green) and anti-DAT (red) that reveals the endogenous protein. (C) Cultured MB dopamine neurons expressing DAT-pH and immunolabeled with anti-GFP (green) and anti-tyrosine hydroxylase (TH) (red) antibodies. From left to right are the cell body, distal axon and the enlarged view of the two cropped areas of the distal axons containing TH-positive (a) and TH-negative (b) axons. (D) Comparison of the expression and co-expression patterns of DAT-pHluorin (DAT-pH) and other versions of tagged-DAT in N2a cells. Top, illustrations of the molecular structure and location of the tag for all DAT reporters. Representative images selected from N=3 repeats. (E) ^3^H-DA uptake for N2a cells, N2a cells expressing DAT-pHluorin, SH-SY5Y cells and N2a cells expressing YFP-DAT. N=3 replicates. (F) Bar graphs of normalized ^3^H-DA uptake for SH-SY5Y cells, N2a cells and N2a cells expressing DAT-pHluorin in response to cocaine (10 μM). N=6 replicates.

When expressed in cultured MB neurons, we found that DAT-pH exhibited homogenous cell body expression similar to endogenous DAT, and it was recognized by an anti-DAT antibody (**Fig. 1B**). DAT-pH was targeted to express in distal axons in both DAergic and non-DAergic neurons in the culture (**Fig. 1C**). In the past, fluorescent proteins such as YFP have been tagged to the N-terminus of DAT to track DAT trafficking in heterologous cells. We next examined the expression of DAT-pH in N2a cells by comparing to other tagged/fluorescent DAT reporters (**Fig. 1D**). Unlike the N-terminally tagged DATs, which often exhibit supraphysiological levels of plasma membrane localization (Li et al. 2004; Sorkina et al. 2009), DAT-pH expression was predominantly intracellular, but exhibited perfect colocalization when coexpressed with DAT-HA (**Fig. 1D**). To determine if a fraction of DAT-pH was indeed targeted to express on the plasma membrane we performed the ^3^H-DA uptake assay. N2a cells transfected with DAT-pH exhibited a near 40-fold increase in ^3^H-DA uptake. Although it was only a fraction of the ^3^H-DA uptake in YFP-DAT expressing cells, DAT-pH in N2a cells performed similarly compared to the endogenous transporter activity measured in SH-SY5Y cells (**Fig. 1E**). A 60-minute cocaine exposure inhibited DA uptake for N2a cells expressing DAT-pH by 90%, similar to the cocaine’s effect on the endogenous transporter in SH-SY5Y cells (**Fig. 1F**). These data further suggested the low-abundance plasma membrane localization of DAT-pH, and that pHluorin insertion did not block the essential substrate binding sites on DAT.

As a reporter gene that was designed to retain endogenous functions, an important concern is the potential unregulated expression that leads to altered intracellular signaling. An earlier study by Ryan and colleagues showed that only 1∼2 copies of vGlut1-pHluorin (equivalent to a 10∼20% overexpression) expression on the synaptic vesicle was sufficient to serve as a working reporter (Balaji and Ryan 2007). Using immunofluorescence labelling of DAT, we found that DA neurons expressing DAT-pH exhibited similar levels of overall DAT expression compared to the untransfected neurons (**Supplemental Fig. 1A-B**). We then analyzed DAT surface expression in transfected and untransfected neurons using a fluorescent cocaine analogue, JHC1-64 (Cha et al. 2005; Eriksen et al. 2009; Guthrie et al. 2020) following a validation study conducted in N2a cells (**Supplemental Fig. 1C-D**). We showed that JHC1-64 can bind to DAT-pH in non-DAergic neurons (**Supplemental Fig. 1E**). Expressing DAT-pH did not lead to an increase in JHC dye binding either at the cell body or the axons (**Supplemental Fig. 1F, G**). The reduced fluorescence at the axons could be due to reduced binding affinity for DAT-pH.

We next examined other functional properties of DAT-pH as a viable reporter. Perfusion of a pH 5.5 MES (2-(N-morpholino)ethanesulfonic acid) solution quenched the DAT-pH fluorescence while a pH 7.4 NH_4_Cl solution revealed its peak fluorescence at the cell body (**Fig. 2A**). To determine if pHluorin insertion in the functional DAT molecule maintains its original pH sensitivity (estimated pKa ∼ 7.1 in synaptopHluorin (Sankaranarayanan et al. 2000), we measured the pKa of DAT-pH in the axons of MB neurons by sequential perfusion of a series of pH-fixed solutions following a previously reported protocol (Sankaranarayanan et al. 2000) (**Fig. 2B-C**). We found the best fitted pKa to be 7.0 for DAT-pH in MB neurons at rest, and that cocaine (10 μM for 4 days) treatment did not alter the pH sensitivity of the reporter **(Fig. 2D**). Moreover, we examined if DAT-pH exhibits agonist-regulated trafficking. DAT has been shown in earlier studies to internalize upon PKC activation (Li et al. 2004; Miranda et al. 2005), and insulin was able to increase DAT surface expression via activation of the PI3K pathway (Carvelli et al. 2002). Consistently, DAT-pH exhibited a time-locked insulin-induced increase in fluorescence and a PMA (phorbol myristate acetate)-induced decrease in fluorescence (**Fig. 2E**), suggesting that the new reporter is responsive to PI3K and PKC regulated trafficking. We then examined psychostimulants including AMPH and cocaine. A brief AMPH (10 μM) perfusion led to a robust decrease of the fluorescence, which returned to baseline upon washout (**Fig. 2F left**). Interestingly, cocaine (10 μM) perfusion also induced a transient reduction of the DAT-pH signal, suggesting DAT internalization, which was followed by fast reversal and an increase of fluorescence within a few minutes (**Fig. 2F right**).

**Figure 2.**
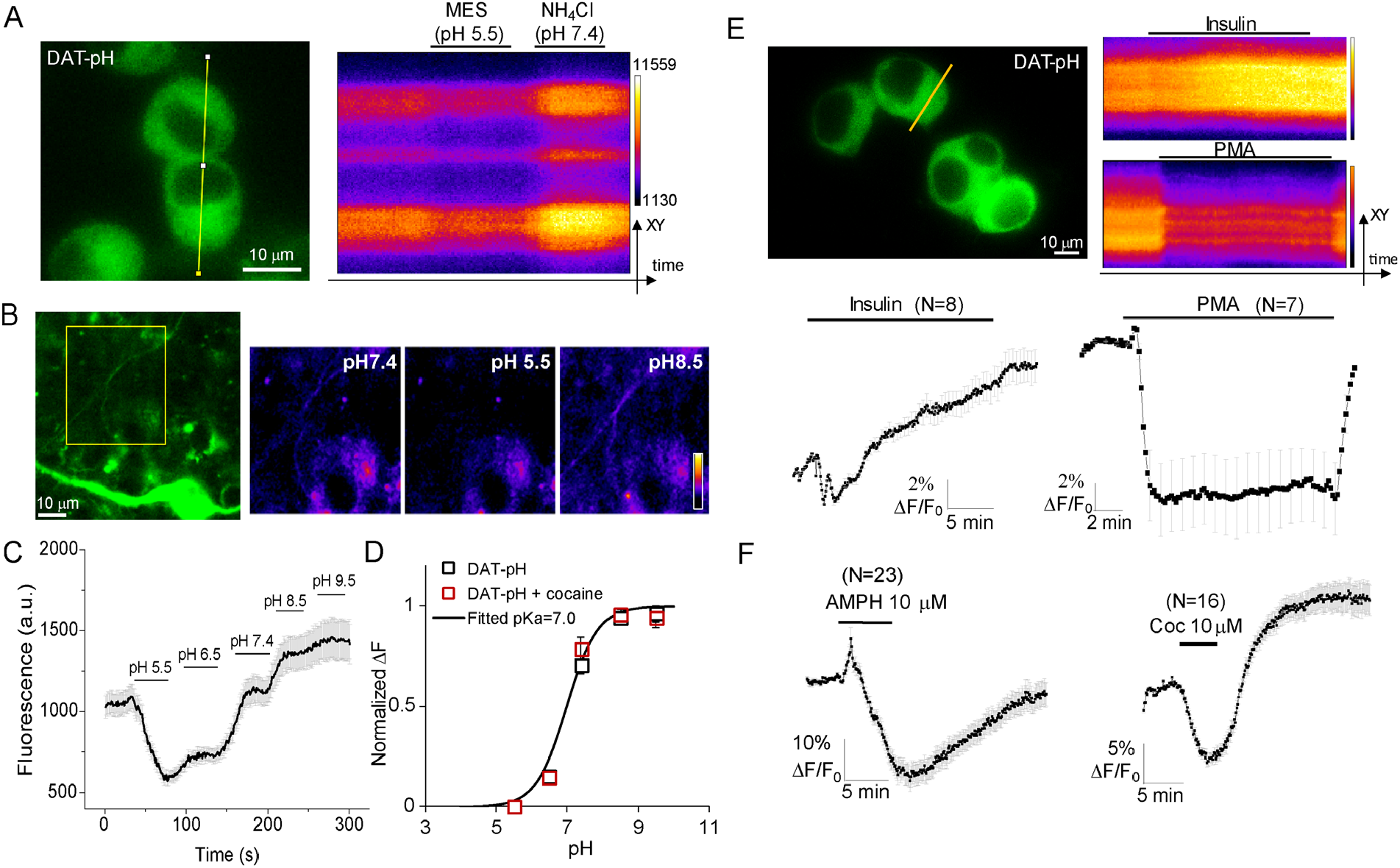
DAT-pHluorin exhibits reliable pH sensitivity and ligand regulated trafficking. (A) N2a cells expressing DAT-pH were perfused with a pH 5.5 MES solution followed by a pH 7.4 NH_4_Cl solution. Right, kymographs were generated from the yellow line of the image on the left. (B) Representative MB neuron expressing DAT-pH and axons (yellow boxed region) were selected for analyzing the pKa. Image intensities were adjusted to visualize the axon. Cell body fluorescence was not saturated at the camera. Right panels are the representative axon with color coding for DAT-pH immunofluorescence taken at different time points during a sequential perfusion of pH-fixed solutions shown in C. (C) Representative DAT-pH signal when perfused with MES buffered saline at pH5.5 and 5.6, NH_4_Cl solution at 7.4, and Bicine buffered saline at pH8.5 and 9.5. Error bars are from standard errors of 31 regions of interests along the axon of one neuron. (D) Summary of normalized changes in DAT-pH responses in (C) from 4 axons at rest (black symbols) and 5 axons after the repeated cocaine treatment (red symbols). Black line represents the fitted change in fluorescence assuming pKa=7.0. (E) N2a cells expressing DAT-pH and DAT-HA were perfused with Tyrode’s solution containing insulin (1 μM) or PMA (1 μM) in separate imaging trials. Kymographs were generated from the yellow lines in the top left panel and the corresponding fluorescence signals of the cells were presented at the bottom. N= cell number, which is indicated next to the trace. Data is represented as mean ± S.E.M. Similar responses were verified in three separate experiments. (F) Averaged responses from N2a cells expressing DAT-pH and DAT-HA perfused with Tyrode’s solution containing amphetamine/AMPH (10 μM, 5 min) or cocaine/coc (10 μM, 5 min). N=cell number. Data is represented as mean ± S.E.M from two independent cultures.

Taken together, our analyses using complimentary methods suggest that DAT-pH is a viable, and likely superior optical reporter for analyzing DAT trafficking in neurons.

### Majority of axonal DAT is on the plasma membrane

We next transfected DAT-pH in cultured mouse midbrain neurons. By bath perfusing MES (pH 5.5) and NH_4_Cl (pH 7.4) solutions, we were able to extract an MES response that represents the surface pool of DAT and an NH_4_Cl response that represents the intracellular DAT (**Fig. 3A-B, supplemental Fig. 2A**). In the cell body and proximal dendrites of the neuron, both surface and intracellular DATs were observed (**Fig. 3A-C**). Interestingly, the intracellular DAT (NH_4_Cl response) in the majority of the axons (identified by their substantially thinner appearance) was either absent or significantly smaller compared to that from the dendrites and cell body (**Fig. 3B-C**). In parallel, the axonal MES response is substantially larger compared to the cell bodies and dendrites (**Fig. 3B, Supplemental Fig. 2B**). To exclude the possibility of quenching intracellular DAT by MES (**Supplemental Fig. 2C**), we performed surface labeling of pHluorin using an anti-GFP antibody without permeabilizing the cell (**Supplemental Fig. 2C-D**). Our analysis revealed the highest ratio for the surface to intracellular labeling of pHluorin in the axons and the lowest ratio at the cell body (**Supplemental Fig. 2E**), further suggesting a negligible contribution from the membrane permeability concerns of MES. To determine if DAT-pH yields different responses in DAergic versus non-DAergic neurons from the MB culture, we performed *post hoc* analysis based on tyrosine hydroxylase (TH) immunoreactivity of the neuron analyzed. Across four independent batches of the MB cultures, we did not observe significant differences in either the MES or NH_4_Cl responses (**Supplemental Fig. 3A**).

**Figure 3.**
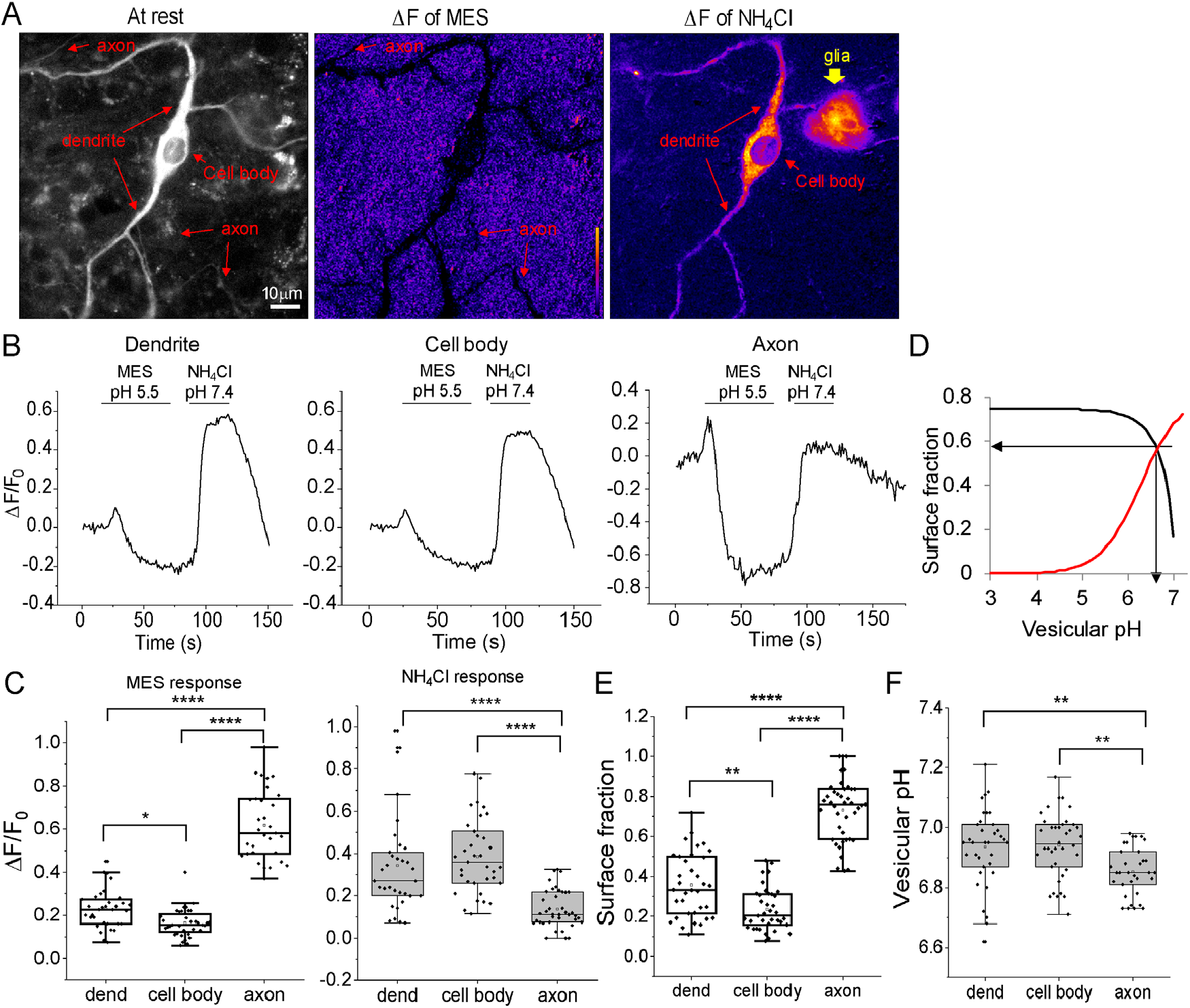
Majority of axonal DAT is on the plasma membrane. (A) Representative imaging field containing a cultured midbrain neuron and a neighboring glia expressing DAT-pHluorin. From left to right: fluorescent image at rest in grey scale, ΔF image of the negative response to the MES solution perfusion and ΔF image of the positive response to the NH_4_Cl solution perfusion. Arrows point to the cell body, neurites and the glial cell, which were visible at different stages. (B) Representative DAT-pHluorin signals measured from the dendrite, cell body and axon during a sequential MES and NH_4_Cl perfusion experiment. (C) Box and whisker charts for the ΔF/F_0_ of the MES and NH_4_Cl responses measured at different compartments of the midbrain neuron. Each data point represents an averaged response from one cell across multiple trials. N=33 for dendrites, N=36 for cell bodies and N=31 for axons. F=159.67, p=2.06E-31, one-way ANOVA for MES responses. ****p=0.0000 comparing axons vs. dendrites and axons vs. cell bodies; *p=0.039 comparing dendrites vs. cell bodies, Tukey’s *post hoc* following one-way ANOVA. F=22.56, p=7.29E-9, one-way ANOVA for NH_4_Cl responses. ****p=4.84E-6 comparing axons vs. dendrites; ****p=1.48E-8 axons vs. cell bodies; p=0.55 comparing dendrites vs. cell bodies, Tukey’s *post hoc* following one-way ANOVA. Data from four batches of cultures. (D) Illustration for the calculation of surface fraction and vesicular pH based on the ΔF/F_0_ measurements in obtained in (B) using the Henderson-Hasselbalch equation (see methods). (E-F) Box and whisker charts summarizing the surface fraction (E) and vesicular pH (F) of DAT-pH in different cellular compartments. F=131.72, p=6.37E-30, one-way ANOVA for surface fraction. ****p=0.0000 comparing axons vs. dendrites and axons vs. cell bodies; **p=0.0012 comparing dendrites vs. cell bodies, Tukey’s *post hoc* following one-way ANOVA. F=7.14, p=0.0013, one-way ANOVA for vesicular pH. **p=0.0059 comparing axons vs. dendrites; **p=0.0025 comparing axons vs. cell bodies; p=0.98 comparing dendrites vs. cell bodies, Tukey’s *post hoc* following one-way ANOVA.

The dynamic range of the DAT-pH response is determined by the acidity of the intracellular structure that it is delivered to. To more accurately determine the surface DAT fraction in each neuronal compartment and the vesicular pH of the DAT-pH in that compartment, we used the Henderson-Hasselbalch equation with the fitted DAT-pH pKa of 7.0 (**Fig. 2B-D**), following previously published protocols (**Fig. 3D**) (Balaji and Ryan 2007; Ashrafi et al. 2017).We report that neuronal cell body and dendrites contain 23.49 ± 1.75% (N=37) and 35.64 ± 2.68% (N=35) of surface DAT, respectively, while axonal DATs are primarily localized to the plasma membrane (73.1 ± 2.38%, N=41) (**Fig. 3E**). On average, DAT-containing vesicles are modestly but significantly more acidic in the axons (pH=6.85 ± 0.14) compared to those in the cell body (pH=6.94 ± 0.17) and dendrites (pH=6.94 ± 0.22) (**Fig 3F**).

### Varicosities are distinct axonal sites that contain DAT in acidic intracellular organelles

The relative distribution of intracellular versus plasma membrane DAT at rest in different subcellular compartments of the neuron is reflective of constitutive DAT recycling. In all axonal structures examined, a fraction of these exhibited a relatively large NH_4_Cl response (**Supplemental Fig. 2B**), suggesting that parts of the axons might be subjected to active DAT internalization. However, it remains unclear whether DAT is internalized at random positions along the axon, or if there are designated structures that support DAT turnover. To address this question, we examined three different structures in the axon: 1) synaptic boutons, which are active sites for synaptic vesicle recycling and exhibit punctate staining for synapsin (**Fig. 4A**); 2) axonal shafts, which exhibit diffuse synapsin staining; as well as 3) varicosities that often exhibit DAT-pH outlining at rest (**Fig. 4B**) but weak or diffuse synapsin staining. These varicosities, although not observed in all axons, are not uncommon for MB neurons. They have been previously reported to be potential axonal branching points (Matteoli et al. 2004) and are presently viewed to be highly relevant in neural injury and neurodegeneration (Gu 2021). We found that only varicosities but not axonal shafts or boutons exhibited a robust NH_4_Cl response (**Fig. 4B-C**). No significant difference was observed for the MES responses (**Fig. 4C**). We further calculated the surface DAT fraction using the Henderson-Hasselbalch equation and found that axonal shafts and boutons exhibited a similar ∼75% plasma membrane fraction of DAT and similar acidity for DAT containing vesicles (**Fig. 4D-E**). No significant difference was observed between TH+ and TH-neurons (**Supplemental Fig. 3B**). Varicosities, however, exhibited only ∼20% plasma membrane DAT (**Fig. 4D**) and DAT-containing vesicles were substantially more acidic with an average pH of 6.31 ± 0.14 (**Fig. 4E**). Thus, our data suggests that the constitutive recycling of DAT is similar between axonal shafts and synaptic boutons, while large varicosities stand out as unique structures potentially involved in DAT endocytosis and degradative processing.

**Figure 4.**
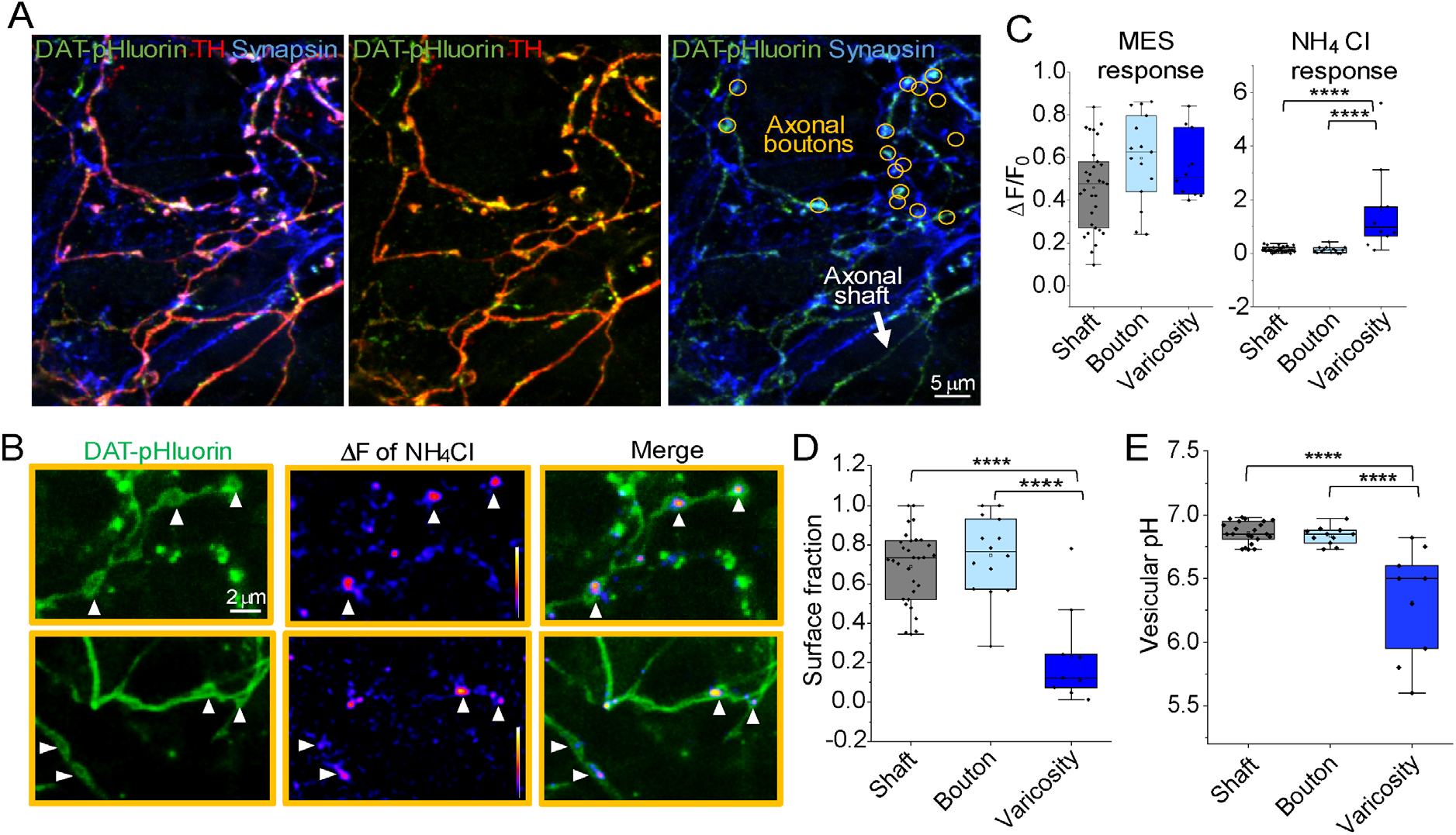
Varicosities contain a large portion of acidic intracellular DAT. (A) Representative immunofluorescence images of the axons expressing DAT-pH labeled with anti-GFP, anti-TH and anti-synapsin1/2. Orange circles indicate synapsin-enriched axonal boutons and the white arrow points to an axonal shaft with light synapsin staining. (B) Representative axonal segments expressing DAT-pH and responded robustly to NH_4_Cl perfusion. From left to right: DAT-pH axon at rest (left), the ΔF image of the NH_4_Cl response of the axon (middle) and the overlay of the two images (right). The arrowheads point to varicosities with DAT-pH contours and NH_4_Cl-responding centers. (C) Box and whisker charts summarizing the MES and NH_4_Cl responses for axonal shaft (N=31), synapsin boutons (N=15) and large varicosities (N=10). F=2.90, p=0.064, one-way ANOVA for MES responses. F=18.07, p=1.10E-6, one-way ANOVA for NH_4_Cl responses. ****p=1.33E-6 comparing varicosity vs. shaft, ****p=1.38E-5 comparing varicosity vs. bouton, p=1.00 comparing bouton vs. shaft, Tukey’s *post hoc* test following one-way ANOVA. (D-E) Calculated surface fraction (D) and DAT-containing vesicular pH (E) based on the Henderson and Hasselbalch equation using the measurements in C. Data from four batches of cultures. F=21.59, p=1.62E-7, one-way ANOVA for surface fraction. ****p=4.86E-7 comparing varicosity vs. shaft, ****p=5.34E-7 comparing varicosity vs. bouton, p=0.66 comparing bouton vs. shaft, Tukey’s *post hoc* test following one-way ANOVA. F=23.63, p=1.90E-7, one-way ANOVA for vesicular pH. ****p=1.90E-7 comparing varicosity vs. shaft, ****p=6.05E-6 comparing varicosity vs. bouton, p=0.94 comparing bouton vs. shaft, Tukey’s *post hoc* test following one-way ANOVA.

### Cocaine exposure elicits a redistribution of DAT to the axonal plasma membrane

Cocaine has been shown to not only inhibit DAT’s reuptake activity but also to increase DAT surface expression (Saunders et al. 2000; Little et al. 2002; Daws et al. 2002; Siciliano et al. 2018), or to prevent DAT from internalization (Underhill et al. 2014; Siciliano et al. 2018; Sorkina et al. 2021). The precise manner by which cocaine changes DAT surface expression/availability, especially in neurons, is not clear. An increase in DAT binding sites were reported at the striatal/synaptic regions of primates after chronic cocaine use (Letchworth et al. 2001; Mash et al. 2002). Whereas, in rodent models of substance use disorder, voltammetry studies have suggested reduced DA uptake, hence, less surface DAT in the nucleus accumbens (NAc) (Calipari et al. 2014; Ferris et al. 2012).

To better understand cocaine-induced DAT trafficking in neurons, and to specifically pinpoint the changes of DAT surface expression in various subcellular compartments of the neuron, we expressed DAT-pH in cultured midbrain neurons and subjected the cell culture to either a 1-day or a 4-day cocaine (10 μM) treatment. We then compared surface DAT fraction and DAT-containing vesicular pH from cocaine treated cells to cells without cocaine treatment from the same batch. Interestingly, DAT surface fraction remained relatively steady at the cell bodies and dendrites (**Fig. 5B**). The axons exhibited a significant increase in surface DAT by day 4 (**Fig. 5B**), which is consistent with the gradual reduction of the NH_4_Cl response (**Fig. 5A**) and a significant alkalization of the DAT-containing vesicles (**Fig. 5C**).

**Figure 5.**
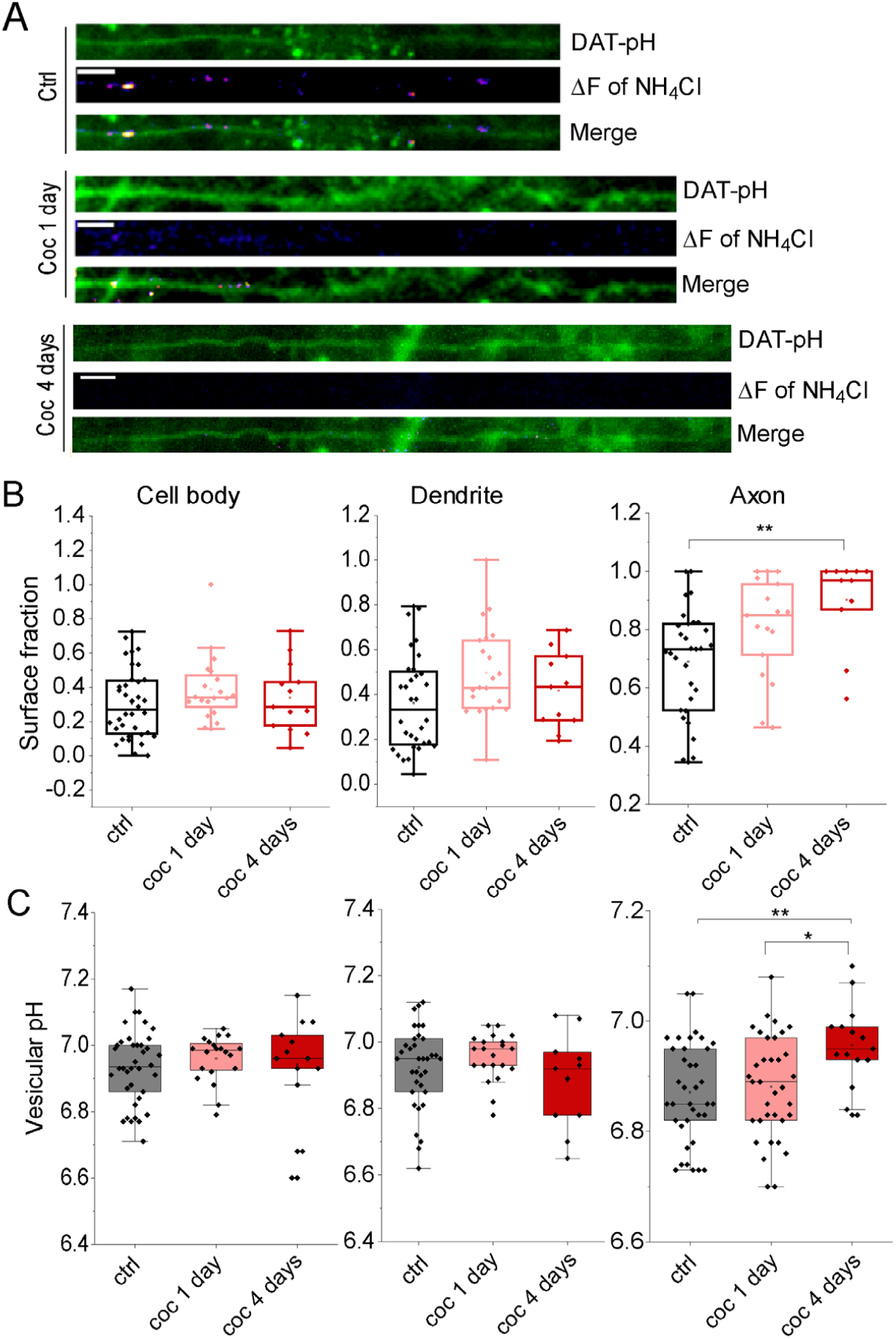
Cocaine exposure results in an increase in surface DAT fraction in the axon. A) Representative NH_4_Cl responses (ΔF of NH_4_Cl) at the axon in cells without exposure to cocaine (control/ctrl), those that have been exposed to cocaine for 24 hours (coc 1 day) and those that have been exposed to cocaine for 4 days (coc 4 days). The baseline DAT-pH images were shown above the NH_4_Cl ΔF images and the overlay images (merge) were shown at the bottom. Scale: 5 μm. (B-C) Box and whisker plots for surface fraction (B) and vesicular pH (C) at the cell body, dendrites and axons in cocaine treated and non-treated neurons. Cell body: ctrl N=32, coc 1 day N=20, coc 4 days N=13, F=1.17, p=0.32 for surface fraction and F=0.34, p=0.71 for vesicular pH, one-way ANOVA. Dendrite: ctrl N=32, coc 1 day N=21, coc 4 days N=11, F=2.78, p=0.070 for surface fraction and F=1.50, p=0.23 for vesicular pH, one-way ANOVA. Axon: ctrl N=31, coc 1 day N=17, coc 4 days N=11, F=6.73, p=0.0024 for surface fraction, one-way ANOVA. **p=0.0030 comparing ctrl vs. coc 4 days, p=0.075 comparing ctrl vs. coc 1 day, p=0.35 comparing coc 1day vs. coc 4 days, Tukey’s *post hoc* following one-way ANOVA. F=5.19, p=0.0075 for vesicular pH, one-day ANOVA. **p=0.0066 comparing ctrl vs. coc 4 days, *p=0.021 comparing ctrl vs. coc 1 day, p=0.88 comparing coc 1day vs. coc 4 days, Tukey’s *post hoc* following one-way ANOVA. Data from six batches of cultures for control, four batches of cultures for coc 1 day and three batches of cultures for coc 4 days.

### The cocaine-induced DAT trafficking is regulated by synaptojanin1

To further investigate the mechanisms underlying cocaine-regulated DAT trafficking, we examined DAT-pH signals in response to a shorter cocaine treatment. Similar to the N2a cells, neuronal cell bodies and axons exhibited a cocaine-induced reduction of DAT-pH fluorescence, which recovered and reversed to an increase in DAT-pH fluorescence within 5 minutes following cocaine removal (**Fig. 6A-B**). Interestingly, in neurons carrying a heterozygous mutation of *Synj1* (*Synj1+/-*), a larger deflection was observed (**Fig. 6A-B**), which could suggest increased endocytosis of DAT or delivery of DAT to more acidic intracellular structures. Nonetheless, the slightly altered internalization phase did not impact the steady-state surface delivery of DAT-pH at the cell body (**Fig. 6A**) but completely abolished this process in the axons of *Synj1+/-* neurons (**Fig. 6B**). *Synj1* encodes a phosphoinositide phosphatase, synaptojanin1 (synj1) which has been well-characterized for its role in clathrin-mediated synaptic vesicle (SV) recycling (de Heuvel et al. 1997; Verstreken et al. 2003; Mani et al. 2007; Milosevic et al. 2011; Watanabe et al. 2018). Synj1 deletion results in an accumulation of clathrin-coated vesicles within the synaptic terminal (Cremona et al. 1999; Harris et al. 2000; Verstreken et al. 2003). Our data demonstrates the first evidence that Synj1 also regulates DAT trafficking in the axons.

**Figure 6.**
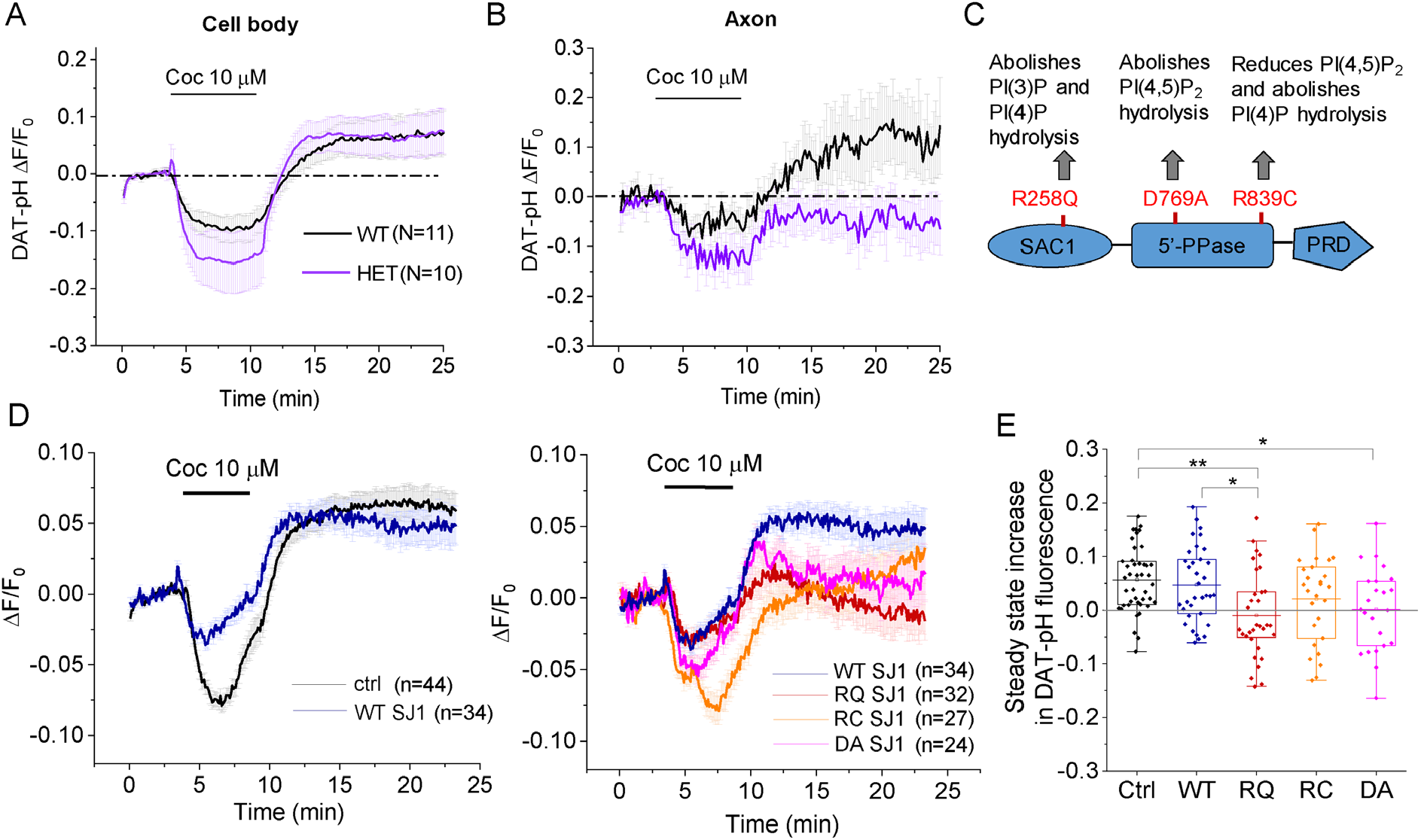
The cocaine-induced DAT trafficking is regulated by Synj1. (A-B) Averaged DAT-pH signals in response to an acute perfusion of cocaine measured from the cell body (A) or axon shafts (B) of the littermate MB neurons. Data is represented as mean ± S.E.M. N=cell number from three independent batches of cultures. (C) Synj1 domain structure, mutants used in the study and the previously reported impact of these mutations on phosphoinositide metabolism. (D) Averaged DAT-pH signals in response to an acute perfusion of cocaine measured from control (ctrl) N2a cells or those transfected with WT hSJ1 or hSJ1 mutants. Left, overlay of the responses from control cells (black) and cells expressing WT SJ1 (navy). Right, overlay of the responses from WT and mutant SJ1 transfected cells. Data is represented as mean ± S.E.M. from two independent batches of cultures. N=cell number. (E) Box-whisker charts for the average of the final 3 min of the DAT-pH responses, which was denoted as steady state increase across all test groups. F=5.21, p=5.78E-4, one-way ANOVA. **p=0.0013 comparing ctrl vs. RQ, *p=0.015 comparing WT vs. RQ, *p=0.033 comparing ctrl vs. DA, p=0.63 comparing WT vs. RC, p=0.14 comparing WT vs. DA, Tukey’s *post hoc* analysis following one-way ANOVA.

We next sought to determine the domain function of Synj1 involved in this regulation by expressing mutant human SYNJ1 (hSJ1) in N2a cells. In our previous work, we have shown that the Parkinson’s disease (PD)-linked R258Q mutation in the SAC1 phosphatase domain abolishes the PI(3)P and PI(4)P hydrolysis activities of the enzyme without affecting the primary 5’-phosphase (5’-PPase) activity; however, the PD-linked R839C mutation reduces the PI(4,5)P_2_ hydrolysis (by 60%) and abolishes PI(4)P hydrolysis (**Fig. 6C**)(Pan et al. 2020). Another mutation in the 5’-PPase domain, hD769A/mD730A, which was previously reported to abolish the PI(4,5)P_2_ hydrolysis (Mani et al. 2007) was also included in this analysis to perform a comprehensive dissection of the lipid signaling pathways. We found that wildtype (WT) SJ1 expression did not influence the surface delivery of DAT despite a smaller internalization response (**Fig. 6D**). Both the RC and DA mutations interfered with DAT surface delivery, however, the RQ mutation in the SAC1 domain exhibited the most statistically significant impairment compared to the WT SJ1 (**Fig. 6D-E**). Thus, our work using a novel DAT optical reporter suggested an essential role of the Synj1’s SAC1 activity in cocaine-regulated DAT trafficking.

## Discussion

DAT trafficking is an integral part of DAT function and bears tremendous importance in DA-related disorders. However, our understanding on neuronal DAT trafficking is limited primarily due to the lack of research tools that are capable of revealing DAT trafficking events in small structures such as the axons. Here, we report the engineering of a novel DAT reporter, DAT-pH. We show that DAT-pH senses pH with a stable pKa of 7.0, while exhibiting DA uptake, cocaine sensitivity and regulated trafficking. We show that in cultured MB neurons, DAT-pH exhibit 25% of surface expression at the cell body but 75% in the axons. We further identified varicosities as potential hotspots for axonal DAT internalization and degradation. Finally, we report a novel role of Synj1, especially its SAC1 phosphatase, in assisting cocaine-induced DAT trafficking to the axonal surface.

As a novel reporter for DAT trafficking, DAT-pH offers multiple advantages: 1) DAT-pH is capable of revealing DAT localization or constitutive recycling in small neuronal structures that otherwise requires electron microscopy (EM) or super resolution imaging analysis. We report a compartmentalized expression pattern of DAT in neurons, which is highly similar to previous EM analyses in rodent brains (Hersch et al. 1997; Nirenberg et al. 1996; Block et al. 2015). These results lend credibility to visualizing actual DAT trafficking events in neurons. 2) DAT-pH is a superior tool for revealing the dynamic changes of DAT in real-time for cell lines and neurons. Fluorescent DAT ligands, which have also been reported to label and track plasma membrane DAT (Eriksen et al. 2009; Guthrie et al. 2020), often exhibit competitive binding with cocaine and native DA. Using DAT-pH we report for the first time the dynamic internalization and surface delivery of DAT upon cocaine exposure, which is regulated by a phosphoinositol phosphatase, Synj1. It is also worth noting that while N2a cells initiated the reverse trafficking of DAT within the 5 minutes of cocaine exposure, neurons did so after cocaine removal. These kinetic differences between a neuron and a heterologous cell, although not yet understood, would not be possible with conventional tools. 3) As a genetically encoded reporter, DAT-pH has the potential to be used *in vivo* or in combination with another optical sensors, such as RdLight1 (Patriarchi et al. 2020) that measures DA. The DAT is an essential molecule for shaping DA transients in the brain, however, little is known about how trafficking of DAT is regulated by DA or other psychostimulants; and whether/how the translocation of DAT in turn modifies DA transients in the brain. For example, the somatodendritic DAT has been shown to not only mediate DA uptake but also participates in reverse transport of DA (Falkenburger, Barstow, and Mintz 2001; Opazo, Schulz, and Falkenburger 2010; Rice and Patel 2015; Sulzer, Cragg, and Rice 2016). Thus, real-time measurement of DAT surface expression and DA transients may advance our understanding of DA signaling in an unprecedented way. There are, however, a few caveats to be noted: First, DAT-pH measures pH rather than direct surface expression, which brings richness to the data (such as the DAT-containing vesicular pH) but also caveats when interpreting a singular measurement in a time-lapse study. Second, DAT-pH reports the relative surface fraction and does not contain information on the overall abundance of DAT. This could bring limitations when considering lateral diffusion as an important form of DAT trafficking in the axon (Bagalkot et al. 2021; Eriksen et al. 2009). Indeed, we showed that the cocaine-induced increase in surface DAT at the cell body (**Fig. 6A**) was not measurable after 24 hours (**Fig. 5B**), suggesting potential lateral diffusion. Future endeavors in developing ratiometric reporters for DAT would circumvent such concerns. Lastly, the expression of the current DAT-pH in neuronal axons are relatively week, which limits the signal/noise for axonal DAT trafficking. Further optimization of the reporter design is required to expand the types of applications for DAT-pH.

Nonetheless, the DAT-pH reporter and its future derivatives are timely tools for investigators across the fields to understand DAT dysregulation in substance use disorders, affective disorders, and Parkinsonian neurodegeneration. Using DAT-pH we revealed a previously uncharacterized mode of axonal DAT trafficking especially in response to cocaine exposure. Interestingly, WT SJ1 overexpression resulted in a less robust internalization but didn’t affect the steady state DAT surface expression. The RQ SJ1 induced a similar internalization as the WT SJ1 but failed to deliver DAT to the surface. Thus, the internalization phase that we observed appeared to be independent from DAT surface delivery. We are yet to understand the biological significance and the molecular mechanisms underlying the cocaine-induced DAT internalization. However, we are the first to show that the cocaine-induced surface expression of DAT is regulated by Synj1. Mutations in both phosphatases of *SYNJ1* have been implicated in early onset Parkinsonism (Krebs et al. 2013; Quadri et al. 2013; Kirola et al. 2016; Taghavi et al. 2018). Remarkably, genetic variants of *SYNJ1* was also associated with bipolar disorder (Horschitz et al. 2005; Saito et al. 2001; Kato, Kakiuchi, and Iwamoto 2007). The lack of ligand-induced DAT trafficking in the axons of *Synj1+/-* neurons may contribute to the altered DA signaling in the pathogenesis of these disorders, which we will investigated in the future. We envision that similar analyses can be carried out to assess other disease risk genes involved in PD, substance use disorders and psychosis.

Taken together, using a novel pH-sensitive DAT reporter our study demonstrated previously uncharacterized biology of DAT trafficking in different parts of the neuron. We suggest immense potential for this novel reporter in revealing greater molecular details of DAT trafficking and DA signaling in physiological states as well as in disease pathogenesis.

## Materials and Methods

### Key reagents and RRIDs

| Reagents | RRID |
| --- | --- |
| YFP-hDAT | Addgene_90228 |
| RFP-HA-DAT | Addgene_90265 |
| SH-SY5Y cells | CVCL_0019 |
| Neuro-2a (N2a) cells | CVCL_0470 |
| C57BL/6J mice | IMSR_JAX:000664 |
| Chicken anti-GFP | AB_2534023 |
| Guinea pig anti-synapsin1/2 | AB_1106784 |
| Mouse anti-TH | AB_477569 |
| Rabbit anti-TH | AB_10077691 |
| Rat anti-DAT | AB_2190413 |

#### Constructs and cloning

Human DAT cDNA was amplified from YFP-hDAT (Addgene, 90228) and pHluorin was amplified from vGlut1-pHluorin (a gift from Timothy A. Ryan and pHluorin is an original asset of James E. Rothman). The hDAT-pHluorin cDNA was inserted in the modified pCAGGS backbone (cut out from the Eevee-iAkt-pm sensor (Miura, Matsuda, and Aoki 2014), a gift from Kazuhiro Aoki) using the ECoR-I and Xba-I restriction sites. The plasmid contains a CAT (combination of chicken beta-actin promoter with CMV enhancer) promotor and a KRas CAAX sequence at the C-terminus for improvement in plasma membrane targeting (Michaelson et al. 2005). An Age-I restriction site was inserted first by two-step PCR amplification using primers at the desired position of hDAT, followed by enzyme digestion and pHluorin insertion. The same linkers in vGlut1-pHluorin were used (TGSTSGGSGGTGG and SGGTGGSGGTGGSGGTG) on both ends of pHluorin in DAT-pH. The correct orientation of the pHluorin was confirmed by sequencing. DAT-HA was amplified from RFP-HA-DAT (Addgene, 90265) using ECoR-I and Xba-I sites and subcloned into the same backbone as the DAT-pH. All primers were synthesized by IDT custom DNA oligo services and all subcloning enzymes and buffers were purchased from New England Biolabs (NEB). DH5α (Thermo Fisher, 18265017) or XL-10 Gold (Agilent Technology, 200314) competent cells were used for plasmid amplification.

The following constructs were purchased directly from Addgene: pEYFP-C1-DAT (90228), tagRFP-C1-DAT-HA (90265) and pCAG-ECFP (32597). CFP-DAT was subcloned using the following primers: forward-5’-gctagcgctaccggtcaagaattcgccacc-3’ and reverse-5’-aattcactcctcaggcgaattctgcagtcg-3’ to replace EYFP using the Age-I and EcoR-I enzymatic sites.

#### Animals

C57BL/6J mice were originally purchased from the Jax lab. The *Synj1+/-* mice (Cremona et al. 1999) were gifted by the Pietro De Camilli laboratory at Yale University. *Synj1+/-* mice were crossed to the C57BL/6J mice to generate littermate pups for MB cultures. Mice were housed in the pathogen-free barrier facility at the Rutgers Robert Wood Johnson Medical School SPH vivarium. Handling procedures were in accordance with the National Institutes of Health guidelines approved by the Institutional Animal Care and Use Committee (IACUC).

#### Cell culture and transfection

N2a cells were cultured and passaged following an ATCC suggested protocol using culture media containing DMEM (Thermo Fisher, 11965118), 10% fetal bovine serum (FBS) (Atlantic Biologicals, S11550), and 5% 10 U/mL Penicillin-Streptomycin (Thermo Fisher, 15140122). Cells were trypsinized using 0.05% Trypsin-EDTA (Gibco, 25300-054) and plated at 30% confluency. Transfection was carried out following a company suggested protocol the day after plating using Lipofectamine™ 3000 (Thermo Fisher, L3000015). The transfection mixture was left in the medium until the day of imaging (typically within 24-48 hours).

SH-SY5Y cells were cultured and passaged according to the ATCC suggested protocol using DMEM/F-12(1:1) culture media (Gibco, 11320-033) containing 10% FBS (Atlantic Biologicals, S11550), and 5%10 U/mL Penicillin-Streptomycin (Thermo Fisher, 15140122). For experiments, cells were trypsinized using 0.05% Trypsin-EDTA and plated at 50% confluency. Transfection was performed as previously described above.

Midbrain cultures (Pan and Ryan 2012; Pan et al. 2017) were prepared as described previously. Ventral midbrains (containing both VTA and SN) were dissected from P0 mouse pups and digested using papain (Worthington, LK003178) in a 35 °C water bath with gentle stirring and constant oxygenation. Midbrain neurons were then seeded within a cloning cylinder on the #1.5 cover glasses pretreated with Poly-L-ornithine (Sigma, P3655). Cells were plated at a density of 199,000 cells / cm^2^ and grown in the Neurobasal-A based medium supplemented with GDNF (10 ng/mL, EMD Millipore, GF030). Lipofectamine™ 2000 (Thermo Fisher, 11668019) was used for transfection at DIV (days in vitro) 7 following a company suggested protocol. The DNA-lipofectamine mixture was washed out after 45 min incubation at 37 °C and the growth medium was replaced with a fresh medium supplemented with an antimitotic agent, ARA-C (Sigma-Aldrich, C6645). For cocaine experiments, cells were treated with 10 μM cocaine (Sigma-Aldrich, C5776) for 24 hours (1 day) or refreshed daily for 4 consecutive days. Treatment typically starts on DIV (days in vitro) 12-13 and imaging experiments were performed between DIV 13 and DIV 17.

#### Radioactive DA uptake assay

Radioactive DA uptake assay was performed following previous published protocols (Aggarwal and Mortensen 2017). N2a cells and SH-SY5Y cells were seeded in triplicates in the 24-well plate two days before the uptake assay. Cells typically reach 80% confluency on day of testing. For cocaine inhibition, cells were treated with 10 μM cocaine (Sigma, C5776) for 50 min in the cell culture incubator prior to the uptake assay based on published results to achieve maximum inhibition (Sun et al. 2017). ^3^H-DA (Dihydroxyphenylethylamine (Dopamine), 3,4-[Ring-2,5,6-3H]-, 250µCi (9.25MBq)) was purchased from PerkinElmer (NET673250UC). 50 nM ^3^H-DA was diluted in the uptake buffer containing 130 mM NaCl, 1.3 mM KCl, 10 mM HEPES, 1.2 mM MgSO_4_, 1.2 mM KH_2_PO4, 2.2 mM CaCl_2_, 10 mM glucose as well as 50 mM L(+) ascorbic acid (Millipore, AX1775-3) and 50 mM pargyline (Sigma Aldrich, P8013.) from freshly prepared stock before the experiment. 10 μM cocaine was added when testing cocaine inhibition. The pH of the buffer was adjusted to 7.4. Cells were incubated with the ^3^H-DA at room temperature for 10 min before washing with an additional uptake buffer without ^3^H-DA. Scintillation fluid was added to the wells immediately and transferred to the tubes. The radioactive signal was read by the scintillation counter (Beckman, LS 6500). For cocaine inhibition analysis, duplicate cultures were made for Western blot analysis and all uptake counts were normalized to the total DAT proteins in the culture.

#### Live imaging and data processing

For live cell imaging, cells on cover glass were mounted on a custom-made laminar-flow chamber with constant gravity perfusion (at a rate of ∼0.2-0.3 mL/min) of a Tyrode’s salt solution containing 119 mM NaCl, 2.5 mM KCl, 2 mM CaCl2, 2 mM MgCl2, 25 mM HEPES, 30 mM Glucose, 10 μM 6-cyano-7-nitroquinoxaline-2,3-dione (CNQX), and 50 μM D, L-2-amino-5-phosphonovaleric acid (AP-5) and buffered to pH 7.40. The NH_4_Cl medium contains: 50 mM NH_4_Cl, 70 mM NaCl, 2.5 mM KCl, 2 mM CaCl_2_, 2 mM MgCl_2_, 25 mM HEPES, 30 mM Glucose, 10 μM CNQX, and 50 μM AP-5, buffered to pH 7.40. The MES medium contains: 25 mM MES, 70 mM NaCl, 2.5 mM KCl, 2 mM CaCl_2_, 2 mM MgCl_2_, 30 mM Glucose, 10 μM CNQX, and 50 μM AP-5, buffered to pH 5.50. Perfusion of Tyrode’s solution is regulated by Valvelink 8.2 and the NH_4_Cl or MES solutions were perfused by pipettes. All chemicals were purchased from Sigma-Aldrich. Temperature was clamped at ∼31.0 °C on the objective throughout the experiment. Images were acquired using a highly sensitive, back-illuminated EM-CCD camera (iXon+ Model DU-897E-BV, Andor Corp., CT, USA). Nikon Ti-2 wide-field microscope is modified with Spectra-X (Lumencor) as the light source for shuttered illumination. pHluorin fluorescence excitation and collection were using a Nikon PLan APO 60X 1.40 NA objective using 525/50m emission filter and 495LP dichroic filters (Chroma, 49002).

Images were sampled at 1 Hz using the Elements software and analyzed in ImageJ. Cell body responses were measured by regions of interests (ROIs) drawn by the freehand tool based on the ΔF image of the NH_4_Cl response. Dendritic responses were measured using the 8×8 pixels (2 μM) circular ROIs based on the ΔF image of the NH_4_Cl response. Axonal responses were measured using the 6×6 pixels (1.5 μM) circular ROIs based on the basal fluorescence image and the ΔF image of the MES response. For varicosity of a large size, several circular ROIs might be placed to cover the area. The average response of all ROIs from one of the three above compartments (cell body, dendrite or axon) was extracted using the Time Series Analyzer plugin. All responses were background subtracted followed by baseline normalization in OriginLabs. Typically, two or three trials of sequential perfusions were carried out with 5-10 min spacing for each imaging field and the averaged peak response was measured as the MES or the NH_4_Cl response of the cell.

#### DAT-pH pKa measurements

Measurements were made at midbrain neuron axons due to the highest surface fraction and least interference with endogenous buffers. For the pH 5.50 and pH 5.60 solutions we used 25 mM MES (pKa 6.1) (Millipore-Sigma, M3671) to replace HEPES in Tyrode’s. For the pH 8.50 and pH 9.50 solutions we used 25 mM BICINE (pKa 8.3) (Millipore-Sigma, B3876) to replace HEPES in Tyrode’s. All solutions were perfused by hand pipetting during a period of 5 min imaging. Typically, data from each axon is averaged from two sequential perfusion experiments. DAT-pH fluorescence changes were normalized to the peak ΔF.

#### Calculation for surface fraction and vesicular pH

The calculation of surface fraction and vesicular pH was performed following previous published methods (Mitchell and Ryan 2004). At any given pH (pH_i_), the deprotonated DAT-pHluorin molecules determines the fluorescence. Based on the Henderson-Hasselbalch equation, the fluorescence change during perfusion of a pH 7.4 membrane-permeable NH_4_Cl solution is determined by the following function assuming pKa = 7.0:

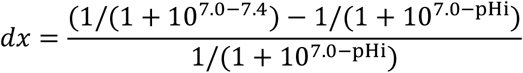

The surface fraction as a function of pH_i_ is then plotted by

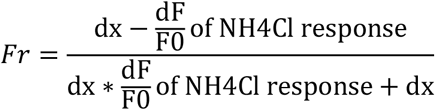

During perfusion of a pH 5.5 MES solution, the ratio of surface to internal DAT-pH can be calculated as the following

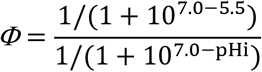

The surface fraction as a function of pH_i_ is then plotted by

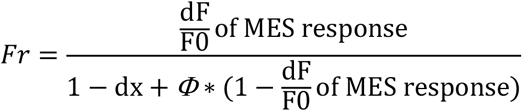

The surface fraction and vesicular pH are determined at the point where two functions intersect.

#### JHC1-64 dye staining

Staining was performed following previously published protocols (Eriksen et al. 2009; Guthrie et al. 2020). Briefly, midbrain cultures were washed three times in ice old JHC-buffer containing: 25 mM HEPES, pH 7.4, with 130 mM NaCl, 5.4 mM KCl, 1.2 mM CaCl_2_, 1.2 mM MgSO_4_, 1 mM L-ascorbic acid, 5 mM D-glucose. Cell culture was then incubated with 50 nM JHC1-64 dye diluted in JHC buffer at 4°C for 30 minutes. After incubation the dye was washed away three times with ice cold JHC-buffer followed by immediate fixation with 4% PFA at room temperature.

#### Immunofluorescence and data analysis

Immunocytochemistry was performed following previously published procedures (Pan et al. 2017; Pan and Ryan 2012). The following antibodies were used for immunofluorescence: chicken anti-GFP (Thermo Fisher, A-10262, 1:1000). Guinea pig anti-synapsin 1/2 (Synaptic System, 106 004, 1:500), mouse anti-TH (Sigma, T2928, 1:1000), rabbit anti-TH (Novus Biologicals, NB300-109, 1:1000), rat anti-DAT (Millipore MAB369 1:1000), Immunofluorescence was analyzed using a Nikon Ti-2 wide-field microscope or a Nikon CREST spinning disk confocal microscope. The same imaging parameters were set for each batch of culture. Image stacks were taken at different focal planes at 0.9 μm interval to include the whole cell and a maximum projection image was generated for each stack via ImageJ for analysis. All analyses were done manually. Synapsin 1/2-positive boutons were determined based on the bright punctate staining pattern.

#### Controlled substances

Cocaine hydrochloride (Sigma C5776) and D-Amphetamine Hemisulfate Salt (Sigma A5880) were purchased from Sigma through authorized institutional distributors.

## Acknowledgements

We would like to thank Dr. Amy Newman for providing the cocaine analogue, JHC1-64 dye; Drs. Zhiping Pang, Timothy Ryan and Jeffrey Diamond for providing valuable insight for the manuscript; and Dr. Loren Runnels for sharing the scintillation counter. The work is supported by an NINDS R01 grant (NS112390) awarded to P-Y Pan, a Rutgers Brain Health Institute Pilot grant awarded to P-Y Pan and D.J. Barker, and a NINDS diversity supplement for J. Saenz (3R01NS112390-02S1).

## Competing Interests

None.

**Supplementary Figure 1 (Related to Figure 1).**
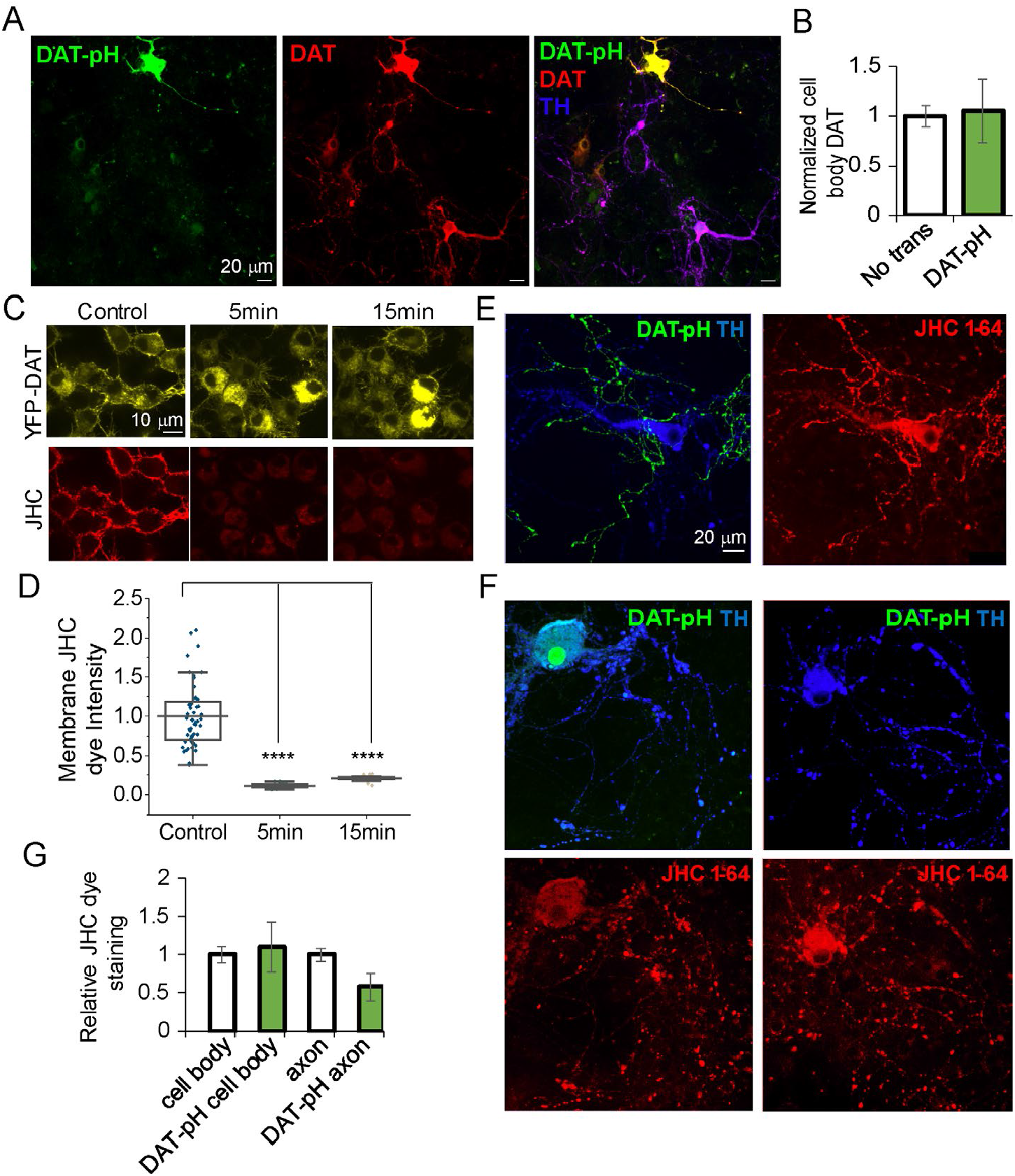
DAT-pH transfection does not lead to overt overexpression of the reporter gene in DA neurons. (A) Representative images of a transfected neuron and a non-transfected DA neuron in the imaging field immunolabeled with anti-GFP (DAT-pH), anti-DAT and anti-TH. (B) Bar graph summarizing the relative total DAT immunofluorescence at the cell body from two batches of cultures. N=26 for non-transfected DA neurons and N=6 for transfected DA neurons. T= -0.19, p=0.85, Student’s *t* test (equal variance). (C-D) Validation of the specificity of the JHC1-64 for plasma membrane DAT using a PMA-induced internalization assay. (C) Representative images of N2a cells transfected with YFP-DAT. After labeling with 50 nM JHC1-64 dye at 4°C for 30 minutes in a staining buffer, following a wash, cells were treated with 1 μM PMA for indicated times, and immediately fixed and imaged using confocal microscopy. (D) Box and whisker charts summarizing the normalized JHC1-64 fluorescence intensity at the plasma membrane after PMA treatment. N=56 for control, N=18 for PMA 5 min, and N=17 for PMA 15 min. F=77.85, p=3.43E-20, one-way ANOVA. ****p=0.0000 for control and 5 min, ****p=0.0000 for control and 15 min, p=0.64 for 5 min and 15 min, Tukey’s *post hoc* following One-way ANOVA. (E-G) JHC1-64 dye analysis for surface expression of DAT in cultured MB neurons. (E) The JHC dye binds to DAT-pH in non-DAergic neurons. (F) Representative DAergic neurons expressing (left) or not expressing DAT-pH (right) stained with the JHC1-64 dye at 4°C to label total surface DAT. (G) Relative JHC 1-64 dye analysis at the cell body and the axons for two batches of the culture containing N=31 non-transfected TH+ neurons and 3 TH+ neurons. For axons, multiple segments were averaged for the same cell. No statistical difference.

**Supplemental Figure 2 (related to Figure 3).**
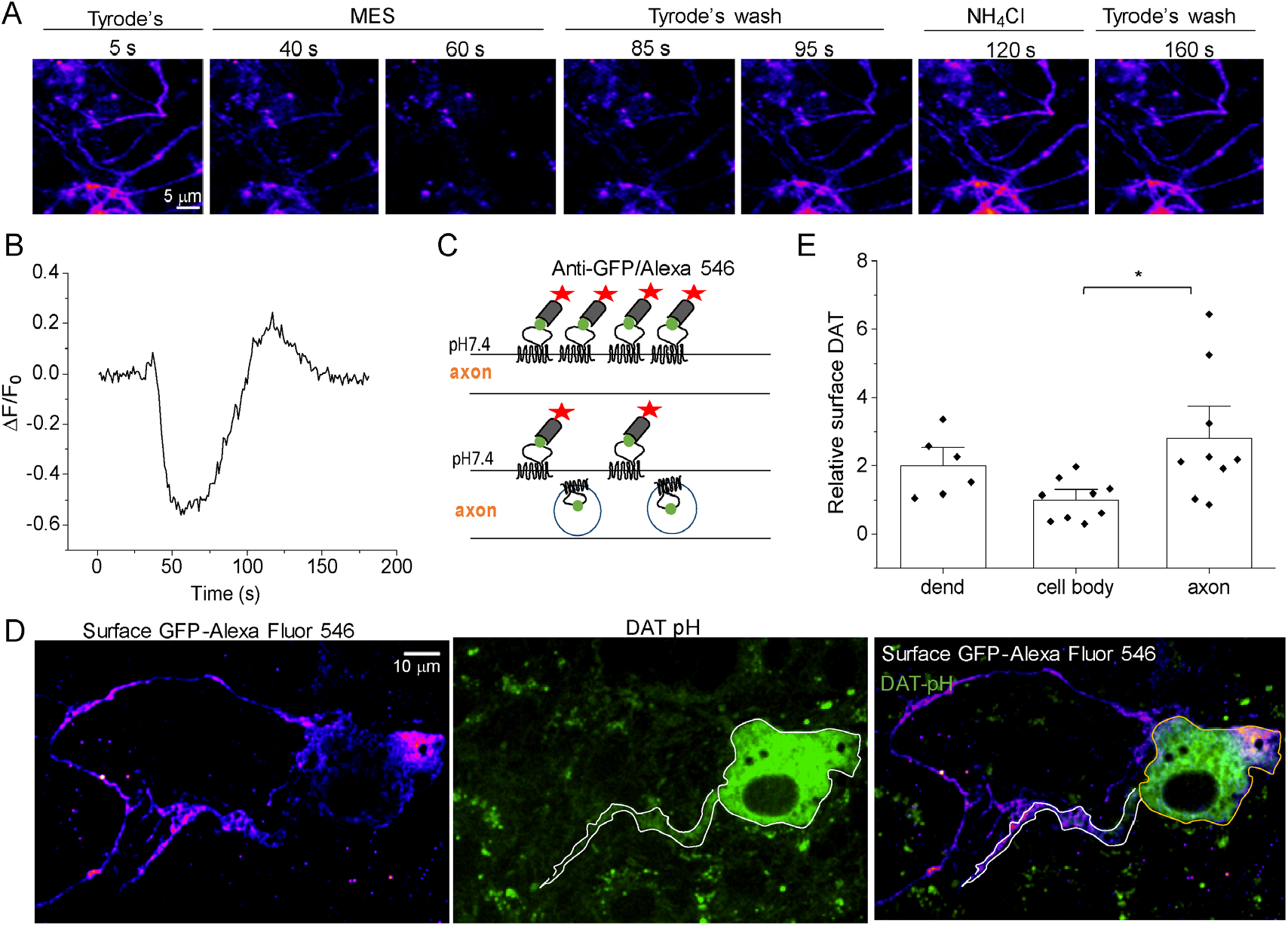
Axonal DAT is largely expressed on the plasma membrane. (A) Time-lapse images of a field of axons expressing DAT-pH perfused with MES and NH_4_Cl The pH 5.5 MES solution almost completely quenched the fluorescence. (B) Representative axonal response from a DAT-pH expressing MB neuron with a relatively large NH_4_Cl response. (C) Illustration for using surface antibody staining to differentiate plasma membrane DAT expression versus intracellular alkaline DAT vesicles. (D) Confocal images of a MB neuron expressing DAT-pHluorin, which was fixed and immunolabeled without permeabilization by Alexa-546 conjugated anti-GFP. Note that axon exhibits the brightest staining compared to the cell body and dendrite. Unlabeled intracellular DAT-pH remained fluorescent, which was only visible at the cell body and dendrite (white outlines). (E) Bar graph for the relative surface DAT (calculated as the Alexa 546 signal/DAT-pHluorin) normalized to the average value of the cell body. Data is presented as mean ± S.E.M. Data from two batches of cultures. N=6 for dendrites, N=9 for cell body and N=9 for axons. F=4.40, p=0.025, one-way ANOVA. *p=0.019 comparing cell body and axons, Tukey’s *post hoc* following one-way ANOVA.

**Supplemental Figure 3 (related to Figure 3 and 4).**
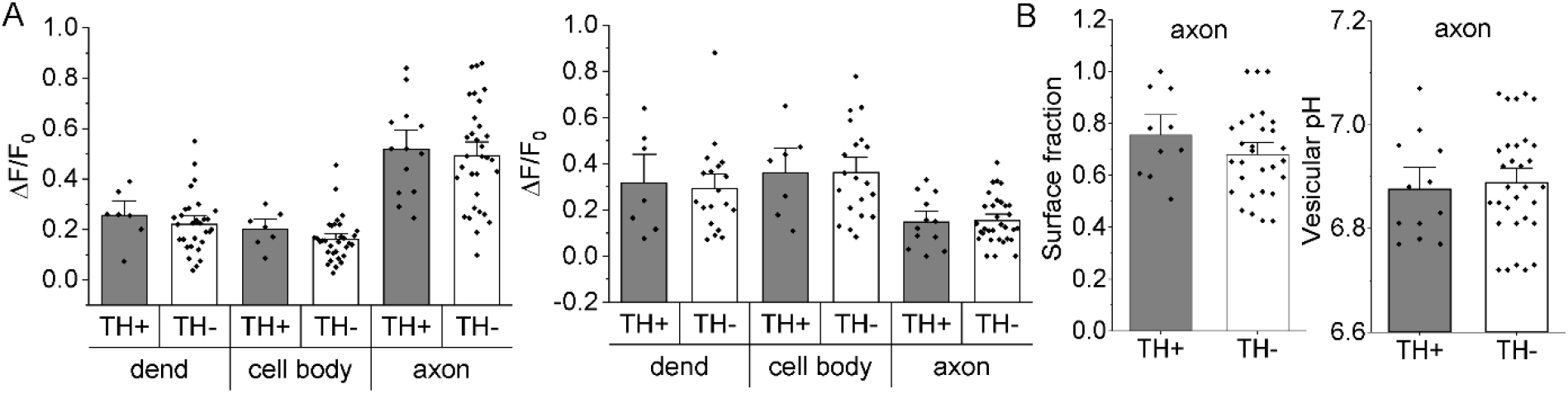
Segregated analysis for MES and NH_4_Cl responses. (A) Bar graphs (mean ± S.E.M.) summarizing *post hoc* analysis for the MES ΔF/F_0_ responses (left) and NH_4_Cl ΔF/F_0_ responses (right) of all batches of neurons based on the TH immunoreactivity at the cell body and neurites examined. N=7 for TH+ dendrites, N=30 for TH-dendrites; N=7 for TH+ cell bodies, N=32 for TH-cell bodies; N=13 for TH+ axons, N=32 for TH-axons. For the MES response, F=42.95, p=1.17E-14 for different compartments, F=0.99, p=0.32 for TH immunoreactivity, two-way ANOVA. For the NH_4_Cl response, F=12.61, p=1.46E-5 for different compartments, F=0.025, p=0.88 for TH immunoreactivity, two-way ANOVA. (B) Calculated surface fraction and DAT-containing vesicular pH for TH+ and TH-axons (including shafts and boutons). N=12 for TH+, N=31 for TH-, T=1.23, p=023, Student’s *t* test (equal variance) for surface fraction; T= -0.35, p=0.73, Student’s *t* test (equal variance) for vesicular pH.

